# Transovarial transmission fuels persistent infections of a core virome in the Chagas disease vector Rhodnius prolixus

**DOI:** 10.1101/2020.10.18.344325

**Authors:** Tarcísio Fontenele de Brito, Vitor Lima Coelho, Maira Arruda Cardoso, Ingrid Alexandre de Abreu Brito, Fides Lea Zenk, Nicola Iovino, Attilio Pane

## Abstract

Triatomine assassin bugs comprise hematophagous insect vectors of *Trypanosoma cruzi*, the causative agent of the Chagas disease. Although the microbiome of these species has been investigated to some extent, only one virus infecting *Triatoma infestans* has been identified to date. Here, we describe for the first time seven (+) single-strand RNA viruses (RpV1-7) infecting *Rhodnius prolixus*, a primary vector of the Chagas disease in Central and South America. We show that the RpVs belong to the Picorna-Calici, Permutotetra and Luteo-Sobemo clades and are vertically transmitted from the mothers to the progeny via transovarial transmission. Consistent with this, all the RpVs, except RpV2 that is related to the entomopathogenic Slow bee paralysis virus, established persistent infections in our *R. prolixus* colony. Furthermore, we show that *R. prolixus* ovaries express 22-nucleotide viral siRNAs (vsiRNAs), but not viral piRNAs, that originate from the processing of dsRNA intermediates during viral replication of the RpVs. Interestingly, the Permutotetra and Luteo-Sobemo viruses display shared pools of visRNAs that might provide the basis for a cross-immunity system. The vsiRNAs are maternally deposited in the eggs, where they likely contribute to reduce the viral load and protect the developing embryos. Our results unveil for the first time a complex core virome in *R. prolixus* and begin to shed light on the RNAi-based antiviral defenses in triatomines.

**Author summary:** *Rhodnius prolixus* is a triatomine insect and a primary vector of *Trypanosoma cruzi*, the etiologic agent of the Chagas disease, in Central and South America. Despite the medical relevance, very little is known about the viruses that infect these so-called assassin bugs. In this study, we show for the first time that triatomines can support the concomitant infection of a variety of RNA viruses belonging to distantly related viral families. Remarkably, we show that the viruses are vertically transmitted from the mothers to the progeny via transovarial transmission. The detection of 22-nucleotide viral small interfering RNAs in mature eggs strongly suggests that RNAi mechanisms contribute to reduce the viral load during oogenesis and embryogenesis in *R. prolixus*, thus safeguarding the development of embryos and nymphs. In agreement with these findings, all the viruses, except one, could establish persistent infections in our colony. Our results substantially expand the knowledge of the virus complexity in triatomine species. This viral toolkit might be harnessed to develop novel insect population control strategies to reduce the diffusion of the Chagas disease.

## Introduction

Triatomine insects (Hemiptera, *Reduviidae, Triatominae*) include hematophagous species that are responsible for the transmission of the Chagas disease, an infectious illness that affects 6-7 million people worldwide [1]. The insect genera *Rhodnius, Triatoma* and *Panstrongylus* harbor well established vectors of *Trypanosoma cruzi*, the etiologic agent of the illness, with *Rhodnius prolixus* being the most important vector in Colombia, Venezuela and other areas in Central and South America. The protozoan is typically transmitted to the human host through the feces and urine of the bug. The Chagas disease is characterized by chronic cardiac, digestive and neurologic alterations, which can culminate in sudden death due to heart failure [1,2]. Approximately 10.000 people die of Chagas disease per year. Although vectorial transmission has only been documented in Central and South America, this disease has already spread to the United States of America, Canada, many European Countries, Australia and Japan due to food-borne transmission, blood transfusions, organ transplantation, laboratory accidents and congenital transmission from mother to child [3–5]. Currently there are no vaccines or cures for the illness and the most efficient methods to reduce its spreading rely on vector control strategies [1].

*R. prolixus* is a hemimetabolous insect and its development proceeds through 5 nymph stages before reaching the winged adult stage [6,7]. Since each molting phase is promoted by a blood meal, the insect is potentially capable of transmitting the disease at any stage of its development. Blood meals are also critical to initiate oogenesis in adult females, where they allow the production of up to 80 eggs per female. Each ovary in this species is formed by 6-8 ovarioles resembling assembly lines (Fig 1A) [6,8–10]. At the anterior region of the ovarioles, a lanceolate structure known as the tropharium harbors both the trophocytes (i.e. nurse cells) and the immature pro-oocytes. The trophocytes are arranged into a syncytium around a central cavity termed the trophic core. At the posterior region of the tropharium, the oocytes resume meiosis and are encapsulated by follicle cells in an orderly fashion to produce the egg chambers [9,11]. In *R. prolixus*, the egg chambers are exclusively formed by the oocyte surrounded by external follicle cells and remain connected to the tropharium via cytoplasmic bridges termed trophic cords [8]. The cords represent transport routes that transfer RNAs, proteins and nutrients produced by the trophocytes from the trophic core onto the growing egg chamber. Oogenesis in *R. prolixus* culminates in the production of eggs encapsulated by a hard chorion that protects the oocyte from dehydration and mechanical stress [12]. However, small pores and micropili along the margin of the operculum allow the exchange of gases and liquids as well as the fertilization process. At the end of the embryonic development, the 1st instar nymphs hatch by displacing the operculum.

**Fig 1.**
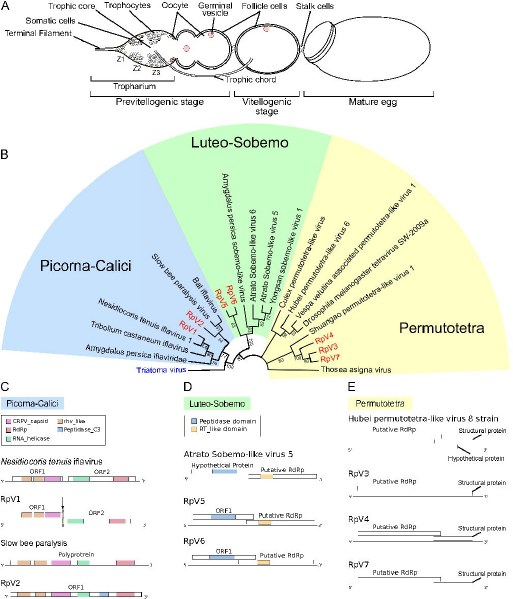
Phylogenetic analysis of the *R. prolixus* viruses. (A) Schematic of an ovariole in *R. prolixus*. Mitotically active trophocytes (i.e. nurse cells) are harbored in Zone 1 of the tropharium, while polyploid trophocytes are present in Zone 2 and 3. The egg chamber in meroistic telotrophic ovaries is formed by the oocyte surrounded by somatic epithelial cells. Trophic cords connect the tropharium to the growing egg chamber and provide transport routes for nutrients and other molecules. (B) Phylogenetic tree of the RpVs constructed with RdRp sequences using Neighbor-Joining method with 1000 bootstrap replicates. Bootstrap values are displayed in tree branches and branches with values less than 50 were collapsed. The RpVs are grouped in three clades: Picorna-Calici, Permutotetra and Luteo-Sobemo. (C) Genome organization of the RpV1 and RpV2 Iflaviruses compared to the closest relatives *Nesidiocoris tenuis* iflavirus and Slow bee paralysis virus respectively. RpV1 displays the 1nt frameshift typical of Iflaviruses, that was not detected in the *N. tenuis* virus. (D) Diagram of the RpV3, RpV4 and RpV7 and their closest relative Hubei permutotetra-like virus 8. (E) Genome organization of RpV5, RpV6 and Atrato Sobemo-like virus 5. rhv-like = Picornavirus/Calicivirus coat protein, CRPV_capsid = Cricket Paralysis Virus capsid protein, RdRp = RNA-dependent RNA Polymerase, RT_like = Reverse Transcriptase_like.

A broad range of arthropod species have been shown to host complex populations of viruses, which can establish harmless persistent infections or affect the behavior, survival and fitness of the host [13–15]. Several members of the Picorna-Calici clade can be either beneficial or pathogenic to their hosts [16]. For instance, the Slow Bee Paralysis Virus (SBPV) transmitted by *Varroa destructor* mites together with other related viruses, like the Kashmir virus and the Deformed wing virus, have been connected to the collapse of honeybee colonies (i.e. Colony Collapse Disorder), which result in severe economic losses [17–19], The *Drosophila* C virus, also a Dicistrovirus like SBPV, was shown to increase the mortality in fruit flies by causing intestinal obstruction and metabolic depression [20–22]. Some arboviruses are well-established agents of zoonosis, like the Dengue, Zika, Chikungunya and Yellow Fever Flaviviruses, which are transmitted by mosquitoes of the *Aedes* family and represent a global health burden [23]. The number of newly discovered arthropod viruses has been exponentially increasing after the development of massive parallel sequencing technologies, but in the majority of the cases, little is known about their biology and ecology as well as their impact on human health and economy [24,25]. It was recently shown that insects host a stable and harmless population of viruses, or core virome, that is vertically transmitted both in laboratory colonies and in nature [26,27].

The host/virus interaction is regulated by the balance between antiviral systems and evasion strategies evolved by the virus. This arms race can culminate in the clearing of the virus from the infected cells, the death of the host or the establishment of innocuous persistent infections. RNA interference is the primary antiviral defense system in insects [28,29]. Typically, double-strand RNA intermediates that are transiently generated during viral replication provide a template for the host Dicer2 (Dcr2) enzyme that cleaves the dsRNA to generate 21-nucleotide (nt) small non-coding RNAs known as viral small interfering RNAs (vsiRNAs). With the help of the R2D2 factor, the vsiRNAs are then loaded into the RNA-induced Silencing Complex or RISC centred on the Argonaute2 (Ago2) protein. The vsiRNAs serve as guides for the RISC to recognize and cleave the viral genomes via the slicing activity of Ago2 [30]. A variety of cellular exonucleases then further degrade the viral sequences and clear the cell from the virus. A growing body of evidence points to a role for another branch of the RNAi phenomena, namely the piRNA pathway, in antiviral defenses in mosquitos [31,32]. In the fruit fly *Drosophila melanogaster*, where the pathway was first discovered, the Piwi-interacting RNAs or piRNAs are 18-30-nt non-coding RNAs that are found in complex with members of the Piwi-clade Argonaute proteins (i.e. PIWIs). In this species, the PIWI/piRNA complexes do not exert antiviral functions, rather they are involved in the silencing of transposable elements and the maintenance of genomic stability. In mosquitos instead, piRNAs originating from viral sequences integrated in the host genome, that is the Endogenous Viral Elements or EVEs, guide the PIWIs to target and eliminate cognate viral genomes [33,34].

The Chagas disease is mainly transmitted by insects belonging to the *Rhodnius, Triatoma* and *Panstrongylus* genera, but more than 150 triatomine species maintain *T. cruzi* infections in the wild and are potential vectors of the disease [3,35]. Despite the medical relevance however, the virome of these insect species has been poorly investigated and to date, only one virus, namely Triatoma virus (TrV), has been described in *Triatoma infestans* [36–38]. Also, RNAi has been extensively used as a tool for functional studies [39,40], but its physiological role in antiviral defense systems remains unexplored in triatomines. In this study, we employed stage-specific *de novo* transcriptome assembly and small RNA profiling to identify novel viruses and investigate the antiviral systems during *R. prolixus* oogenesis. We show for the first time that R. p*rolixus* harbors a complex core virome that is vertically transferred from the mothers to the offspring via transovarial transmission. Furthermore, we find that viral siRNAs, but not viral piRNAs, are produced during oogenesis and likely contribute to protect *R. prolixus* germ cells and early embryos and to establish persistent viral infections. Our findings shed light on the complexity of the virome in triatomine species and their antiviral defense systems and provide a new toolkit for the development of vector population control strategies.

## Results

### Identification of seven novel (+) ssRNA viruses in *R. prolixus* ovaries

We have recently described RNA-Seq transcriptomes from previtellogenic stages of oogenesis (PVS) and mature chorionated eggs (Egg) of *R. prolixus* [39]. In this study, we further interrogated those datasets and discovered that at least 39,340 (0.1%), 46,413 (0.2%), 51,905 (0.2%) and 139,648 (0.6%) reads were related to viral sequences in our RNA-Seq libraries PVS_1, PVS_2, Egg_1 and Egg_2, respectively. *De novo* transcript assembly initially provided 8 contigs in PVS and 7 in Egg stages. Alignment of the contigs in PVS with those found in Egg ultimately led us to identify 7 unique viral genomes, which we labeled *Rhodnius prolixus* Virus 1-7 (RpV1-7) (Fig 1). Multiple sequence alignments and NCBI Blast analyses of the putative RNA-directed RNA polymerases (RdRp) or Open Reading Frames (ORFs) allowed us to establish the phylogenetic relationships among the RpVs and closely related viruses from other arthropods (Fig 1B). Te RpVs belong can be grouped in three different clades: Picorna-Calici (RpV1 and RpV2), Permutotetra (RpV3, RpV4 and RpV7) and Luteo-Sobemo (RpV5 and RpV6) (Fig 1B-E) [25]. None of the RpVs shares similarity with TrV, the only known triatomine virus even though TrV also belongs to the Picorna-Calici clade.

Two of the longest contigs correspond to novel viruses of the Iflavirus genus (Picorna-Calici clade). RpV1 is represented by a 9.6 Kilobases (Kb) long contig and displays two main ORFs encoding a 1,248 AA and a 1,632 AA putative proteins sharing 36.99% and 40.88% amino acid (AA) sequence identity the protein encoded by the ORF1 and ORF2 of the *Nesidiocoris tenuis* iFlavirus, respectively [41] (Fig 1B and C). Remarkably, RpV1 is also highly similar to Deformed wing virus (~32% AA identity), a well characterized entomopathogenic virus. ORF2 encodes the putative RdRp enzyme and a RNA helicase, while ORF1 codes for the capsid proteins typical of Picornavirales (rhv-like) and Cricket Paralysis virus (CRPV). RpV1 ORF1 and ORF2 display a 1-nt overlap typical of Iflaviruses, that was not observed in *N. tenuis* virus [41] (Fig 1C). The 5’ and 3’ Untranslated Regions (UTRs) of RpV1 are 759 and 214 respectively. Although the size of RpV1 is close to the typical length of iFlaviruses (i.e. ~10kb), it is possible that our RNA-Seq library preparation protocols failed to fully capture the sequences at the 5’ and 3’ ends. Similar limitations in defining the exact 5’ and 3’ ends might also apply to the other RpVs described in this study. RpV2 also appears to belong to the Iflavirus genus, but its genome encodes a single polyprotein 2,890 AA in length, that displays ~70% AA identity with the polyprotein of the SBPV [42]. Closer relatives of RpV2 in the phylogenetic tree are also the Bat iflavirus, Moku virus and Vespa velutina virus (Fig 1B and C). The longest contig for RpV2 is ~8.9Kb long and the encoded polyprotein is similar to that of the SBPV with RhV-like and CRPV capsid proteins at the N-terminal region and the RdRp enzyme harbored in the C-terminus (Fig 1C).

Three novel viruses appear to belong to the Permutotetra clade. One of the best characterized viruses of this clade is the *Thosea asigna* virus that displays a typical T=4 icosahedral symmetry of the capsid [43]. The contigs for the Permutotetra-like RpV3, RpV4 and RpV7 are ~5Kb long (Fig 1D). The genomes of these RpVs display two partially overlapping ORFs with ORF1 encoding the RdRp and ORF2 the putative capsid and structural proteins. The putative RdRp enzymes of RpV3, RpV4 and RpV7 share ~42% to ~47% AA sequence identity with proteins from the closely related Shuangao and Hubei permutotetra-like viruses (Fig 1B and D). ClustalW multiple sequence alignment of the RpV3, RpV4 and RpV7 genomes revealed extensive nucleotide sequence identity in any of the combinations (~74% between RpV3 and RpV4, 73% between RpV4 and RpV7 and 78% between RpV3 and RpV7) (S1 Table). Permutotetra-like viruses typically display a 40nm diameter non-enveloped capsid.

Finally, RpV5 and RpV6 belong to the Luteo-Sobemo clade. This clade harbors a broad variety of well-characterized plant viruses that are often phytopathogenic and severely affect agriculture [44,45]. Although these viruses cycle between plants and insects, including beetles, aphids and ticks, their interaction with the invertebrate hosts has been poorly investigated. RpV5 and RpV6 display ~41% and ~66% AA sequence identity with Atrato sobemo-like virus 6 respectively (Fig 1A and D). The size of both RpV5 and RpV6 contigs is ~2.8Kb with two overlapping ORFs, whereby ORF1 encodes the Peptidase activity and ORF2 the RdRp enzyme. The alignment of the putative RdRp enzymes encoded by these viruses showed 70% AA sequence identity, while their genomes share very low nucleotide sequence identity (<30%). Members of the Sobemo-like viruses are best characterized in plants, while much less is known about the insect variants.

Our data reveal for the first time that *R. prolixus* ovaries support the concomitant infection of a range of phylogenetically distant (+) single strand RNA viruses.

### Simultaneous RpVs infections are detected during *R. prolixus* oogenesis

The detection of viral sequences in previtellogenic stages of oogenesis as well as in mature unfertilized eggs suggested that the RpVs might be vertically transmitted from the mothers to the offspring. We therefore sought to investigate this process quantitatively by comparing the steady-state RNA levels of each virus in the PVS and Egg (Fig 2). The endogenous *Rp-rp49* gene, that is abundantly expressed in *R. prolixus* oogenesis, is displayed for comparison (Fig 2A and B) [39]. The comparison of the normalized reads over the two stages of oogenesis shows that the RpVs were generally present both in PVS as well as in the chorionated eggs (Fig 2A and B). Despite the variability between the biological replicates, we found that RpV1 was the most abundant virus in *R. prolixus* ovaries at the time that the RNA-Seq libraries were prepared (2015). However, the accumulation of each RpV in a specific stage of oogenesis may vary. For instance, RpV3 RNA levels were ~100 fold higher in Egg compared to the PVS (Fig 2A and B). Conversely, RpV4, RpV5, RpV6 and RpV7 levels were higher in PVS versus Egg, with RpV6 being almost undetectable in chorionated eggs (Fig 2A and B). The differential accumulation of the RpVs during *R. prolixus* oogenesis was also apparent when we looked at the coverage of the read coverage along each viral genome (Fig 2C-E and S1A-D Fig). This analysis also revealed that the RNA-Seq coverage along the RpV1, RpV3 and RpV4 genomes is apparently uneven. Higher coverage is observed at the 3’ end of the RpV1, while this region shows lower coverage in RpV3 and RpV4 (Fig 2D-E). Although the bias might be due to the protocols employed in the current study, we cannot rule out that the uneven profiles might be due to the accumulation of defective viral particles or subgenomic RNAs corresponding to portions of the viral genomes. These data demonstrate that the RpVs can infect both the early and late stages during *R. prolixus* oogenesis.

**Fig 2.**
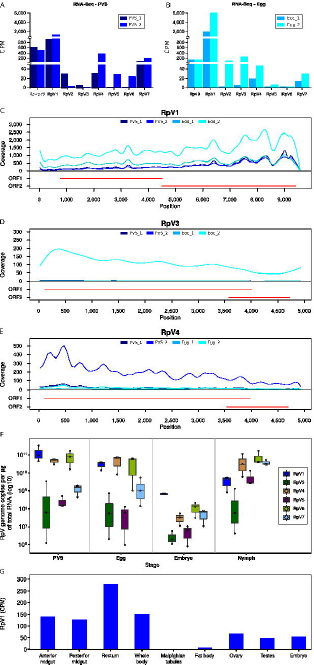
Stage-specific quantification of the RpVs during *R. prolixus* development. (A) Normalized RNA-Seq reads for each virus in previtellogenic stages (PVS) of oogenesis. (B) Normalized RNA-Seq reads for each virus in unfertilized mature eggs (Egg) dissected from the abdomen of adult females. Biological replicates for both stages of oogenesis are scaled by counts per million (CPM). The endogenous *Rp-rp49* gene is also displayed to underscore the abundance of the RpVs in ovarian tissues. Read coverage over the RpV1 (C), RpV3 (D) and RpV4 (E) viral genomes. Red lines indicate the organization of the genome for each virus. (F) Quantification of the RpVs in previtellogenic stages (PVS), mature eggs (Egg), embryos and 1st instar nymphs. RpV2 genome copies were below qRT-PCR detection limits and were not included in the box plot. Y-axis displays the number of viral genome copies per microgram of total RNA (Log10). (G) Quantification of RpV1 in different tissues of *R. prolixus* using available RNA-Seq datasets [46]. Y-axis displays the Counts per Million (CPM).

### RpVs are vertically transmitted and establish systemic infections in *R. prolixus*

All the RpVs are detected in mature eggs dissected from the abdomens of the *R. prolixus* females. This observation prompted us to investigate whether the viruses can actually be passed onto the developing embryos and ultimately to the hatching nymphs. To answer this question, we performed qRT-PCR assays with oligonucleotides specific for each RpV using total RNA samples extracted from PVS, mature eggs, embryos and 1st instar nymphs as templates (Fig 2F). For each stage, we calculated the number of viral genome copies per microgram of total RNA (S2 Table). Compared to 2015, when RpV1 was the most abundant virus in PVS and Egg as per RNAseq, it appears that also RpV4, RpV5 and RpV7 are currently present at levels comparable to RpV1 in these stages of oogenesis (Fig 2F). All the RpVs, except RpV2, are detected also in the developing embryos and 1st instar nymphs (Fig 2F). RpV3 is the least abundant virus in all evaluated developmental stages ranging from 2 × 10^6^to 6 × 10^7^ median number of copies/ug of RNA in embryos and PVS, respectively. Interestingly, the levels of all the RpVs are several orders of magnitude lower in the embryos compared to non-oviposited eggs and 1st instar nymphs (Fig 2F). For instance, RpV1, RpV4 and RpV6 display median levels ranging between 10^10^ and 10^11^ copies/ug of RNA in Egg, but in embryos their levels drop below 10^9^ for RpV1 and 10^8^ for RpV4 and RpV6. In the 1st instar nymphs, the median levels increase again above 10^9^ for RpV1 and 10^10^ for RpV4 and RpV6. RpV2 was below detectable levels in all the assays we performed and thus it was likely lost from our insectarium some time after the transcriptome datasets were produced.

Next, we asked whether the RpVs are able to establish systemic infections and spread to other tissues and organs in *R. prolixus*. Our qRt-PCR and transcriptomic analyses showed that the RpVs, except RpV2, are able to establish persistent infections in this insect. Thus, we interrogated the RNA-Seq datasets published by Ribeiro and collaborators in 2012, who had already noticed viral sequences in their tissue-specific transcriptomes [46]. Despite the small size of each library ranging from 54.2Mb to 240.9Mb, we could clearly detect sequences of RpV1 in the whole body, anterior and posterior midgut, fat bodies, rectum, testes, ovaries, and embryos, but not in the malpighian tubules (Fig 2G). Fewer reads corresponding to RpV2, RpV4 and RpV6 sequences were detected in malpighian tubules and whole body datasets (data not shown).

Collectively, these results show that RpVs can establish persistent and systemic (at least for RpV1) infections in *R. prolixus* via vertical transovarial transmission from the mother to the offspring.

### viral siRNAs for all the RpVs, except RpV2, are detected in early oogenesis and in mature eggs

RNA interference was shown to provide a primary antiviral system in fruit flies and mosquitos [29]. Because our insects seem to support the simultaneous infection and high viral titers of the RpVs, we asked whether a RNAi system might be active in *R. prolixus*. To answer this question, we generated and analysed small RNA datasets from PVS and Egg stages separately. When we mapped the reads against the RpVs, we could identify a total of 113,759 and 139,556 “viral” reads in PVS and Egg, respectively. The length distribution of the small RNAs revealed a clear 22-nt peak for all the viruses except for RpV2 in both stages of oogenesis (Fig 3A and B). As expected, we also did not find small RNAs for TrV that was never detected in our transcriptomes. We then analysed the strand bias of the small RNAs with respect to the viral genomes (Fig 3C and S2 Fig). The small RNA profiles show that both sense and antisense small RNAs for all the RpVs, except RpV2, are expressed in PVS as well as in chorionated eggs. Despite some hotspots, the distribution of the vsiRNAs seems rather uniform along the viral genomes and approximately equal amounts of sense and antisense vsiRNAs are detected for all the RpVs suggesting that they originate from the processing of viral dsRNA replication intermediates (Fig 3C-H and S2 and S3 Figs).

**Fig 3.**
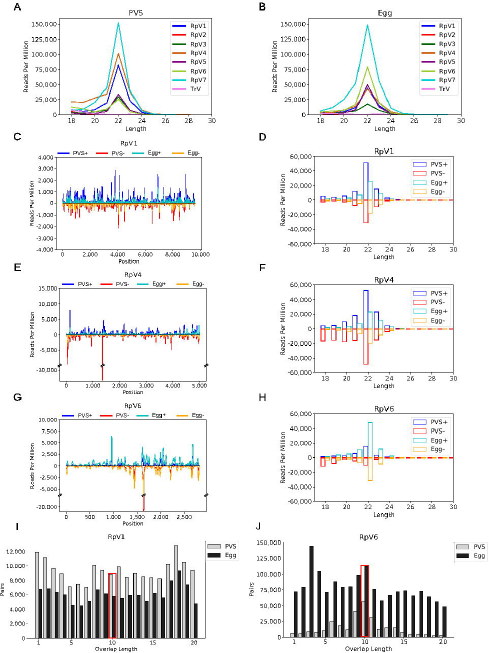
vsiRNA profiling during *R. prolixus* oogenesis. (A) Length distribution of the RpVs small RNAs in previtellogenic stages (PVS) of oogenesis. (B) Length distribution of the viral small RNAs in mature eggs (Egg). In both stages of *R. prolixus* oogenesis, a peak at 22nt is readily detectable for all the RpVs, except for RpV2 and TrV, which did not generate vsiRNA reads above background levels. RpV2 was lost from our insectarium by the time that the small RNA datasets were produced. TrV was never detected in our colony and was used as control for the background small RNA levels. (C-H) vsiRNA profiling along the RpV1, RpV4 and RpV6 genomes and length distribution of the sense and antisense vsiRNAs. Sense and antisense vsiRNAs detected in previtellogenic stages of oogenesis (dark blue = sense, red = antisense) and mature chorionated eggs (light blue = sense, orange = antisense) are displayed for each virus. (I-J) Ping-pong signal for the RpV1 and RpV6 viruses in PVS (white bars) and Egg (grey bars). The red box highlights the position of the typical 10nt overlap between sense and antisense small RNA pairs.

It has been recently proposed that the piRNA pathway might contribute to antiviral defenses in mosquitos [31]. Seminal studies in the fruit fly *D. melanogaster* revealed that the piRNAs range from 23 to 30nt in length and their biogenesis requires the activity of the Piwi-clade Argonaute proteins Aub and Ago3 [47,48]. These enzymes coordinate a feedforward amplification loop termed "ping-pong" mechanism that generates sense and antisense secondary piRNAs with a typical 10nt overlap at their 5'ends. Furthermore, sense and antisense piRNAs are characterized by a Uridine bias at position +1 and an Adenine bias at position +10 respectively. However, when we analysed these features in our RpVs visRNA pools, we did not find strong evidence for a ping-pong amplification mechanism. First, we did not detect viral small RNAs with a 23-30nt size range. Second, we did not find evidence of a ping-pong mechanism (Fig 3I and 3J and S4 Fig). One exception might be represented by RpV6, whose small RNAs display a 10nt overlap enrichment in PVS (Fig 3J). Yet, the RpV6 small RNAs also display enrichments of the 9nt and 11nt overlaps in this stage of oogenesis, that are not generally observed for piRNAs, and the 10nt overlap is not detected in Egg stages. Also, we do not find evidence for the +1 U-bias and +10 A-bias when we analyse the nucleotide frequencies along the viral small RNAs (S5 and S6 Figs).

Although our data strongly point to the absence of viral piRNAs in *R. prolixus* ovaries, we asked whether the viral small RNAs might help identify EVEs in this species. Roughly 2-3% of the PVS and Egg vsiRNAs map both to the viral as well as to the insect genome. However, ~50% of these multi mappers match sequences in protein-coding genes either in intronic (40.22%) or exonic (9,76%) sequences (S3 Table). The remaining vsiRNAs map to intergenic regions in the *R. prolixus* genome, but neither they appear to cluster together nor the cognate genomic sequences are related to the viral genomes. These results demonstrate that the piRNA pathway does not exert a critical role in protecting *R. prolixus* ovaries against RpVs infections.

Intriguingly, we noticed that subsets of visRNAs for the RpV3-7, but not for RpV1 and RpV2, appeared to map to more than one viral genome (Fig 4). Upon closer inspection, we found that the Permutotetra-like RpV3, RpV4 and RpV7 share pools of vsiRNAs in PVS, whereby 217 are common to RpV3 and RpV4, 1553 to RpV3 and RpV7 and 404 to RpV4 and RpV7 (Fig 4A). Remarkably, 717 vsiRNAs are shared among the three viruses (Fig 4A). These findings are supported by the analysis of the Egg datasets, where comparable numbers are observed (Fig 4B). Similarly, the Sobemo-like viruses RpV5 and RpV6 display shared vsiRNAs in PVS (279) and in Egg (762) (Fig 4B). We then asked what regions of the viral genomes produce the shared set of vsiRNAs. To answer this question, we profiled the distribution of the shared vsiRNAs along the viral genomes (Fig 4C and 4D). Surprisingly, both for the Permutotetra-like (Fig 4C) and the Sobemo-like (Fig 4D) RpVs, the shared vsiRNAs mostly map to the 3’ of the genome rather than to sequences encoding the conserved RdRp or structural proteins.

**Fig 4.**
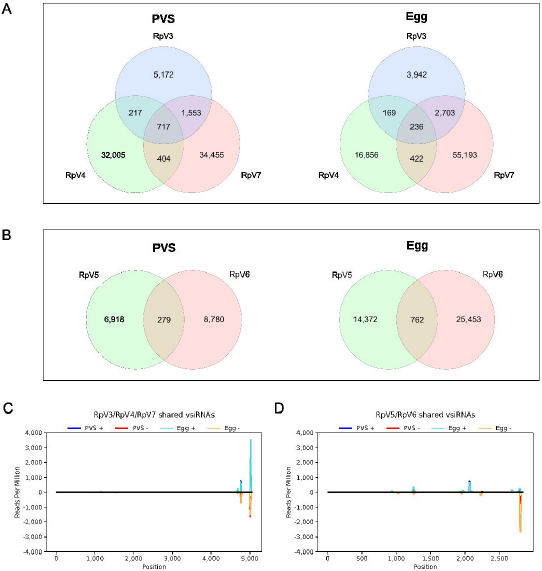
Analysis of the vsiRNA pools shared by the RpVs. (A) Venn diagram showing the vsiRNAs expressed in PVS and Egg stages that are shared by the Permutotetra-like viruses. (B) Venn diagram showing the vsiRNAs expressed in PVS and Egg stages that are shared by the Sobemo-like viruses. Only vsiRNAs with perfect match to the viral genomes were considered in these analyses. (C) Small RNA-Seq coverage of the vsiRNAs simultaneously shared by the Permutotetra-like viruses RpV3, RpV4 and RpV7. The coverage along the consensusgenome (see Methods) is displayed to highlight the accumulation of the shared vsiRNAs in the viral 3'UTR. Similar results were obtained for the RpV4 and RpV7 genomes (S7 Fig). (D) Small RNA-Seq coverage of the vsiRNAs common to the Sobemo-like viruses RpV5 and RpV6. Sense and antisense shared vsiRNAs detected in previtellogenic stages of oogenesis (dark blue = sense, red = antisense) and mature chorionated eggs (light blue = sense, orange = antisense) are displayed.

### Components of the RNA interference mechanisms are conserved in *R. prolixus*

The detection of vsiRNAs in *R. prolixus* ovaries points to the existence of a RNAi machinery that controls the biogenesis and function of these non-coding RNAs. Three branches of RNAi mechanisms have been described in *D. melanogaster* [28]. The canonical RNAi is centered on the Dcr2 and Ago2 proteins, while miRNAs require the Dcr1. and Ago1 proteins. Differently, piRNAs are produced by the activity of the PIWI proteins, Aubergine, Piwi and Ago3 and act in concert with a dedicated set of proteins and enzymes in *D. melanogaster*. A variety of additional factors, including the R2D2 and Loquacious helicases, the Drosha and Pasha nucleases and export factors like Exportin 5 act at different steps in the biogenesis and function of the three classes of small non-coding RNAs. We have previously shown that *R. prolixus* harbors four PIWI genes, with *Rp-piwi2, Rp-piwi3* and *Rp-ago3*, but not *Rp-piwi1*, being expressed both in previtellogenic stages of oogenesis and mature eggs [39]. The analysis of the transcriptomic datasets from chorionated eggs further confirms these findings (Fig 5). More importantly, Blast analyses conducted in VectorBase allowed us to identify the *R. prolixus* orthologs of the main components of the RNAi pathways described in *D. melanogaster* (Fig 5 and S4). Of these, the *Rp-ago2, Rp-piwi2, Rp-r2d2* and *Rp-fmr1* genes display the highest expression levels in ovaries. Except for *Rp-piwi2* that seems to be expressed twice as much in PVS than in Egg, the expression levels of all the genes appear to be comparable in early oogenesis (PVS) and in mature chorionated eggs (Fig 5). Thus, functional RNAi machineries are likely active in early *R. prolixus* oogenesis and are maternally deposited in the mature eggs.

**Fig 5.**
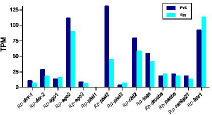
Expression levels of key RNAi genes in *R. prolixus* ovaries. *R. prolixus* putative orthologs of the key factors acting in the *D. melanogaster* miRNA, siRNA and piRNA pathways were identified by Blast analyses using VectorBase, FlyBase and NCBI databases. Expression levels for each gene were computed using the PVS and Egg RNA-Seq datasets. The y-axis displays Transcripts per Million (TPM).

## Discussion

*R. prolixus* is a hematophagous triatomine insect responsible for the transmission of the Chagas disease in Central and Latin American countries. To our knowledge, the composition of the *R. prolixus’* virome is completely unknown and only one virus (i.e. Triatoma virus) is known to infect triatomines, which comprise hundreds of species that are potential vectors of the Chagas disease. Here we substantially expand the virome in these insect species using *R. prolixus* as a model system. We show that *R. prolixus* harbors a complex core virome comprising a range of (+)ssRNA viruses belonging to the Picorna-Calici, Permutotetra and Luteo-Sobemo clades. We demonstrate that RpVs established persistent infections in our colony that were maintained via transovarial transmission from the mother to the offspring. Accordingly, viral genomes can be detected not only in early stages of oogenesis and in mature non-oviposited eggs, but also in developing embryos and in 1st instar nymphs. It is possible that the transport of viral particles through the trophic cords might contribute to the accumulation of the viruses in the developing egg chambers and mature chorionated eggs. Vertical transmission is being extensively studied for medically relevant viruses, like Dengue and Zika virus and has been proposed as a possible explanation for recurrent seasonal epidemics [49–51]. The virus stored in the mosquito eggs, that are resistant to desiccation, might be protected and overcome the unfavorable seasons, when the population of the mosquito vector is reduced. Our results demonstrate that vertical transmission of the RpVs occurs in *R. prolixus* and accounts at least in part for the persistence of the viruses in our insectarium, although additional mechanisms including horizontal transfer via cannibalism e coprophagy might be in place. It will certainly be of great interest to investigate the virome composition and transmission routes in wild caught *R. prolixus*.

Our study also reveals for the first time that RNA interference is active during *R. prolixus* oogenesis and likely protects the developing germline and embryos from viral infections. Indeed, we show that 22nt vsiRNAs for all the viruses, except for RpV2, are detected both in previtellogenic stages of oogenesis as well as in mature eggs. The length of the vsiRNAs in the Hemipteran *R. prolixus* is similar to that observed in some Hymenopteran and Lepidopteran insect species [52]. It seems therefore reasonable to conclude that the RpVs infect the tropharium and early egg chambers, where their dsRNA replication intermediates are converted into vsiRNAs by the *R. prolixus* Dcr2 and associated factors. Both the vsiRNAs and the transcripts encoding the RNAi factors are then loaded into the growing oocyte most likely via the trophic cords, and stored in the mature chorionated eggs. Interestingly, the number of viral genomes for all the RpVs drops by several orders of magnitude in embryos compared to the eggs prior oviposition, but it increases again in the 1st instar nymphs. It is tempting to speculate therefore that maternally provided RNAi machinery might be responsible to reduce the viral levels during embryogenesis in order to safeguard the development of the embryos and nymphs. Conversely, our data demonstrate that the piRNA pathway does not coordinate a prominent antiviral branch during *R. prolixus* oogenesis, even though *Rp-piwi* genes are expressed in the ovaries of this insect species and required for female adult fertility [39].

The observation that the RpVs are not completely eliminated from the ovaries despite an active RNAi machinery is not surprising since it has been shown that the RNAi defenses do not always clear the viruses from the infected cells, rather they seem to reduce the viral burden and avoid the death of the host. This mechanism is thought to sustain persistent viral infections both in plants and animals [53,54] and might account for the fact that most of the RpVs have been maintained in our colony for five years at least, with RpV1 exceeding eight years-long infections. The only exception is represented by the RpV2 virus that was lost from our colony sometime between 2015 and 2017. The closest relative of RpV2 in the phylogenetic tree is the SBPV that is known to cause foreleg paralysis and death in honeybees. It is possible that the replication of RpV2 might have reached critical levels for the survival of the infected insects, whose death eventually interrupted the transmission of the RpV2 in the insectarium. Additional studies will be necessary to determine whether the RpVs are responsible for the periodic weakening and occasional collapse of our colony.

Interestingly, the members of the Permutotetra (RpV3, RpV4 and RpV7) and Luteo-Sobemo (RpV5 and RpV6) clades, display common pools of vsiRNAs, that mostly correspond to sequences in their 3’ end. Hundreds of vsiRNAs can also target any combination of two Permutotetra-like RpVs. These findings seem to point to a cross-immunity mechanism whereby ovarian tissues can use a shared set of visRNAs to prevent or mitigate the replication of distinct, but closely-related viruses. Such a mechanism would also provide an adaptive arm in that the visRNAs for an existing viral infection can be potentially employed to fight a new infection from a phylogenetically related virus. Furthermore, the shared vsiRNA complement might allow the rapid establishment of permanent infections from newly infecting viruses of the same family as well as support the evolution of novel viral variants, while still protecting the host cells from possible damage and death. This hypothesis warrants further investigation in the light of the rapidly increasing viral diversity that is being discovered in arthropods [25].

Interestingly, both enveloped and non-enveloped viral particles were detected in *Trypanosoma cruzi*, the etiologic agent of the illness [55]. Our results suggest that the trypanosome might exchange viruses with the host *R. prolixus*. Given the medical relevance of these organisms, it will be of great interest to thoroughly investigate the interaction between the RpVs, the triatomine insect and the trypanosome as well as the related antiviral defense mechanisms. The employment of metatranscriptomic studies in field-captured triatomines coupled with a thorough analysis of the viral entomopathogenicity will help understand the virome diversity in the triatomine vectors and the possible use of certain viruses in insect population control strategies.

## Materials and Methods

### Rhodnius handling, total RNA extraction and RNA-Seq library preparation

The *Rhodnius prolixus* colony was maintained at 28°C and 75% relative humidity and regularly fed on New Zealand white rabbit blood. Animal handling and experimental protocols were conducted in accordance with the guidelines of the Committee for Evaluation of Animal Use for Research (Universidade Federal do Rio de Janeiro, CAUAP-UFRJ) and the NIH Guide for the Care and Use of Laboratory Animals (ISBN 0-309-05377-3). Protocols were approved by CAUAP-UFRJ under registry #IBQM155/13. Dedicated technicians in the animal facility localized at the Instituto de Bioquímica Médica Leopoldo de Meis (UFRJ) carried out all protocols related to rabbit husbandry under strict guidelines, with supervision of veterinarians to ensure appropriate animal handling. Previtelloegenic stages and mature unfertilized eggs (~30 eggs per biological replicate) were dissected in ice cold 1X PBS from adult females two weeks after the blood meal. Embryos were collected over a period of 5 days after oviposition and roughly 30 embryos were used for each biological replicate. Approximately, 10 1st-instar nymphs per biological replicates were employed. Total RNA was extracted with Trizol Reagents (Life Technologies) as per manufacturer instructions and treated with turbo DNA-free kit (Ambion). Total RNA extracted from unfertilized eggs dissected from femaltes’ abdomens were subjected to paired-end RNA sequencing (RNA-Seq) library production (Illumina) as previously described [39]. RNAseq libraries were generated and sequenced on HiSeq Illumina platforms at Lactad Facility (University of Campinas/Brazil). For small RNA library preparations, 20ug of total RNA was separated on 15M TBE-Urea gels and small RNAs ranging in size 18-30nt were selected as previously described [56]. Library preparation and sequencing were performed at the Max Planck Institute Sequencing Facility with Illumina protocols and NextSeq sequencing platforms.

### PCR and qRT-PCR

Quantitative RT-PCR (qRT-PCR) assays to validate the RNA-Seq datasets and investigate the stage-specific abundance of the viruses were performed as previously described [39,57]. Briefly, 1μg of total RNA for each developmental stage was subjected to genomic DNA removal with the turbo DNA-free kit (Ambion) and retrotranscription with Multiscribe Reverse Transcriptase (ThermoFisher Scientific). Three biological replicates were produced for each assay. The resulting cDNAs were used for qPCR assays with the set of oligonucleotides listed below. Approximately 50ng of cDNA for each sample was mixed with specific oligonucleotides and SYBR green reagent (Life Technologies). qRT-PCR was carried on QuantStudio 7 Flex (ThermoFisher). Analysis of the qRT-PCR results was performed using absolute quantification with standard curves. The standard curve was generated using a 747nt long cDNA corresponding to a portion of the RpV1 virus. The cDNA fragment was amplified by PCR in ovarian cDNAs with oligonucleotides bearing a T7 promoter sequence at the 5’ end as listed in Table 1. The PCR product was subjected to *in vitro* transcription with the T7 Megascript kit (Ambion) followed by DNA removal (Turbo DNA free kit, Ambion). 1μg of the resulting DNA free RNA was converted into cDNA and serial dilutions were used in qRT-PCR assays to produce the standard curve. The number of viral genome copies per microgram of total RNA (Q) for each virus was calculated by *Q* = 10^(*CT–b*)×*m*−1^, where CT is the cycle threshold for a given virus under different conditions, *b* is the intercept-y of the standard curve and *m* is the slope of the standard curve (QIAGEN, 2014). Subsequently, the number of copies was normalized by viral genome size.

### Identification of viral genomes by *de novo* assembly

First we filtered out the host sequences aligning the PVS and Egg RNA-Seq data [39] to the *R. prolixus* reference RproC3 [58] genome using STAR v2.7.2b [59] (--outFilterMultimapNmax 30; --outFilterMismatchNoverLmax 0.1) Unmapped reads were used for *de novo* transcriptome assembly with the Trinity method v2.5.1 [60] with default parameters to identify novel transcripts that were not previously detected due to incomplete sequencing and assembly of the *Rhodnius* genome for each development stage PVS and Egg. *De novo* assembled transcripts were post-processed to filter out potential artefacts based on abundance and quality. First, expression levels of the *de novo* assembled transcripts were quantified by Salmon v1.2.1 [61] and isoforms with TPM < 10 were removed. Next, deduplication of redundant contig sequences was performed by CD-HIT-EST v4.6 [62,63] at a nucleotide identity of 95%. TransRate software [64] used the unmapped reads and the remaining contigs from the previous filtering steps as input to evaluate common assembly errors (chimeras, structural errors, incomplete assembly and base errors) and to produce a diagnostic quality score for each contig, thereby removing possible artefacts. Assembled transcripts remaining from this last step were considered to be good quality transcripts. To identify potential protein-coding transcripts, we compared the assembled contigs against the protein databases of *Rhodnius prolixus* e Swissprot from Uniprot release 2020_01 [65] using BLASTX v2.9 (e-value < 1e-20). We selected the longest sequences with similarity to virus proteins or without hit and blasted against the non-redundant protein database (https://blast.ncbi.nlm.nih.gov).

After the identification of the seven candidate virus sequences, we used the tool rascaf (RnA-seq SCAFfolder) to extend the draft assembly using the RNA-Seq data information [66] (minimum support for connecting two contigs equals to 10 mapped reads). In addition, we performed an all vs all BLASTN alignment v2.10.1 [67] between the PVS and Egg final contigs to identify overlappings that could aid us to extend the viral sequences (default parameters). Alternatively, we performed an independent assembly of the PVS and Egg library reads using the assembler rnaSpades [68] with default parameters. The latter was decisive for the length improvement of RpV2 and RpV7 genomes.

### Virus phylogeny and genome organization

Using the Conserved Domain Database [69], *R. prolixus* viruses were searched in order to identify conserved RNA-dependent RNA polymerase (RdRp) domains in each ORF (Open Reading Frame). The domain regions were then extracted using an in-house script. When RdRp sequences were not readily available in NCBI’s Protein database (https://www.ncbi.nlm.nih.gov/protein), the same methodology was applied to the other viruses: Triatoma Virus (NC_003783), Shuangao permutotetra-like virus 1 (KX883439.1), Culex permutotetra-like virus (LC505019.1), *Thosea asigna* virus (NC_043232), Hubei permutotetra-like virus 8 (KX883453.1), *Vespa velutina* associated permutotetra-like virus 1 (MN565051.1), *Drosophila melanogaster* tetravirus SW-2009a (GQ342964.1), Hubei permutotetra-like virus 6 (NC_033114.1), *Amygdalus persica* iflaviridae (MN823678.1), Slow bee paralysis virus (NC_014137.1), Bat iflavirus (NC_033823.1), *Tribolium castaneum* iflavirus (MG012488.1), *Nesidiocoris tenuis* iflavirus 1 (NC_040675.1), Atrato Sobemo-like virus 6 (MN661101.1), *Amygdalus persica* sobemo-like virus (MN831439.1), Atrato Sobemo-like virus 5 (MN661107.1), Yongsan sobemo-like virus 1 (MH703049.1). Multiple sequence alignment was performed using ClustalW [70]. The phylogenetic tree was constructed with MEGA X [71] using the Neighbor-Joining method [72], bootstrap test with 1000 replicates [73] and Jones-Taylor-Thornton (JTT) matrix [73,74] to compute evolutionary distances.

Gene predictions were performed with the Viral Genome Annotation System (http://cefg.uestc.cn/vgas/) [75] (gene length ≥ 60 nucleotides; ATG, GTG, TTG as start codon). ORF candidates were predicted using NCBI ORFinder (https://www.ncbi.nlm.nih.gov/orffinder/) (ORF length = 75; standard genetic code; ATG and alternative initiation codons as start codons). Protein domains and families were predicted by Conserved Domain Database v3.18 [69] (e-value ≤ 0.05). To compare the RpVs genome organization with known viruses, sequences of the latter were subjected to ORF and protein family prediction methods with the same parameters.

### RNA-Seq bioinformatic analyses

To obtain the RNA-Seq profiles of the virus genomes, we mapped the reads of the PVS and Egg stages [39] against the genome index of the seven RpV genomes using STAR [59] version 2.7.2b (--outFilterMultimapNmax 30; --outFilterMismatchNoverLmax 0.1). Script bamCoverage v3.1.3 [76] produced BigWig read coverage files from the mappings normalized by CPM and bin size equal to 5. Profile plots were visualized using a local instance of the Genome Browser [77].

Expression levels of RNAi components were obtained from the quantification of the RNA-Seq data from pre-vitellogenic stage and unfertilized mature egg [39]. Salmon 1.2.1 [61] was set to produce aggregated gene-level abundance estimates (-g). The transcriptome index was built using k-mer size equal to 19 (-k 19) and the DNA sequences of the improved transcriptome produced by Coelho et al, unpublished.

To assay the presence of our viruses in different *R. prolixus* tissues and organs, we mapped the 454 Roche sequencing data of anterior midgut, posterior midgut, rectum, whole body, malpighian tubule, fat body, ovary, testes and embryo [46] against the genome index of the seven RpV genomes using STARlong [59] version 2.7.2b (--seedPerReadNmax 2000; --outFilterMultimapNmax 30; --outFilterMismatchNoverLmax 0.1).

### SmallRNA-Seq bioinformatic analyses

Raw smallRNA-Seq reads had adapter sequences removed and low quality ends trimmed using TrimGalore! (https://www.bioinformatics.babraham.ac.uk/projects/trim_galore/) with default parameters and filtering out reads with less than 18 nucleotides in length (--length 18), while qualities of sequences were evaluated using FastQC (https://www.bioinformatics.babraham.ac.uk/projects/fastqc/). Genome index for *R. prolixus*, TrV and RpV genomes was made with Bowtie [78] using bowtie-build with default parameters. SmallRNA-Seq reads were then mapped using Bowtie’s “-v –best –strata” mode, allowing up to 3 mismatches. Genome coverage for each virus was performed using Bedtools [79] with genomecov (-d -ibam -strand) and normalized by Reads Per Million method, where the number of reads mapped in each position was multiplied by one million and divided by the number of total reads mapped to each virus. The same normalization method was used for the length distribution of mapped reads in Figs 3A, 3B, 3D, 3F and 3H.

Multi mapped reads that mapped to more than one virus were filtered out and counted. The cons software, part of the EMBOSS package [80], was used to generate consensus sequences with only shared regions between different combinations of RpV genomes (RpV3/4/7, RpV3/4, RpV3/7, RpV4/7 and RpV5/6) in order to identify sites possibly responsible for the production of these small RNAs. SmallRNA-Seq reads were then mapped to these consensus genomes and genome coverage was performed and normalized as described before.

Reads that mapped to *R. prolixus*’ genome were filtered and mapped to RpV genomes. Reads that mapped to both were selected and overlapped with *R. prolixus*’ genomic features present in VectorBase [81] using the intersect software, part of the Bedtools package.

To check if reads mapped to *R. prolixus* viruses displayed any signal of a ping-pong amplification mechanism, reads mapped to the positive strand in each virus were used to search for read pairs mapped in the opposite strand, which displayed a 5’ to 5’ overlap. Overlap between pairs that mapped in more than one place were counted as (1/N1 + 1/N2) / 2, where N1 is the number of different mappings for the read mapped on the positive strand and N2 is the number of different mappings for the read mapped in the negative strand.

## Acknowledgements

We thank Pedro L. de Oliveira, Shree N. Tanneti and Patricia Garcez for critical reading of the manuscript. We are grateful to Helena Araujo, Bernardo Carvalho, Pedro L. de Oliveira and Marcia Cury el Cheikh for the constant help and support. We thank Maria Fernanda Pereira Farias, Kelli Cristina Melquiades Mendes, Graciela Venturi and Luciana da Conceição Soares for invaluable technical support.

## Supporting informations

**S1 Fig. Read coverage and genome organization of Rhodnius prolixus viruses.** (A) RpV2, (B) RpV5, (C) RpV6 and (D) RpV7.

**S2 Fig. Length Distribution of small RNAs along the genome of *R. prolixus* viruses.** (A) RpV2, (B) RpV3, (C) RpV5, (D) RpV7.

**S3 Fig. Length distribution of smalIRNAs mapped in the positive and negative strand of *R. prolixus* viruses.** (A) RpV2, (B) RpV3, (C) RpV5 and (D) RpV7.

**S4 Fig. 5’ to 5’ overlap length distribution of smalIRNAs mapped to *R. prolixus* viruses.** (A) RpV2, (B) RpV3, (C) RpV4, (D) RpV5 and (E) RpV7. Red boxes highlight the 10 nucleotide overlap typically found between piRNA sequences.

**S5 Fig. Nucleotide frequency for each position in vsiRNAs mapped to *R. prolixus* virus 1 to 7 and Triatoma virus in the Previtellogenic stage.**

**S6 Fig. Nucleotide frequency for each position in vsiRNAs mapped to *R. prolixus* virus 1 to 7 and Triatoma virus in the mature egg.**

**S7 Fig. vsiRNAs mapped along shared regions in consensus genomes.** (A) RpV4 and RpV7, (B) RpV3 and RpV7, (C) RpV3 and RpV4.

**S1 Table. Similarity Between R. prolixus viruses.**

**S2 Table. Absolute quantification of RpV.**

**S3 Table. vsiRNAs mapped to Rhodnius and Viruses.**

**S4 Table. R. prolixus RNAi pathway Orthologs.**

**S5 Table. Read abundance per Virus (PVS and Egg).**

**S6 Table. Virus abundance on tissues (454 data).**

**S7 Table. SmalIRNA-seq Read Abundance.**

**S8 Table. Shared smalIRNA-Seq Reads.**

## References

1. WHO. Chagas disease (also known as American trypanosomiasis). [cited 15 Oct 2020]. Available: https://www.who.int/news-room/fact-sheets/detail/chagas-disease-(american-trypanosomiasis)

2. Pérez-Molina JA, Molina I. Chagas disease. The Lancet. 2018. pp. 82–94. doi: 10.1016/s0140-6736(17)31612-4

3. Coura JR, Viñas PA. Chagas disease: a new worldwide challenge. Nature. 2010. pp. S6–S7. doi:10.1038/nature09221

4. Schmunis GA. The globalization of Chagas disease. ISBT Science Series. 2007. pp. 6–11. doi:10.1111/j.1751-2824.2007.00052.x

5. Schmunis GA, Yadon ZE. Chagas disease: A Latin American health problem becoming a world health problem. Acta Tropica. 2010. pp. 14–21. doi: 10.1016/j.actatropica.2009.11.003

6. Nunes-da-Fonseca R, Berni M, Tobias-Santos V, Pane A, Araujo HM. Rhodnius prolixus: From classical physiology to modern developmental biology. Genesis. 2017;55. doi:10.1002/dvg.22995

7. Buxton PA. THE BIOLOGY OF A BLOOD-SUCKING BUG, RHODNIUS PROLIXUS. Transactions of the Royal Entomological Society of London. 2009. pp. 227–256. doi: 10.1111/j.1365-2311.1930.tb00385.x

8. Huebner E. Nurse cell-oocyte interaction in the telotrophic ovarioles of an insect, Rhodnius prolixus. Tissue Cell. 1981; 13: 105–125.

9. Huebner E, Anderson E. A cytological study of the ovary of Rhodnius prolixus. I. The ontogeny of the follicular epithelium. J Morphol. 1972; 136: 459–493.

10. Lutz DA, Huebner E. Development and cellular differentiation of an insect telotrophic ovary (Rhodnius prolixus). Tissue Cell. 1980; 12: 773–794.

11. Pratt GE, Davey KG. The Corpus Allatum and Oogenesis in Rhodnius Prolixus (Stål.). J Exp Biol. 1972;56: 201–214.

12. Beament JW. Memoirs: The Formation and Structure of the Chorion of the Egg in an Hemipteran, Rhodnius prolixus. J Cell Sci. 1946;s2-87: 393–439.

13. Chen YP, Pettis JS, Collins A, Feldlaufer MF. Prevalence and transmission of honeybee viruses. Appl Environ Microbiol. 2006;72: 606–611.

14. Fullaondo A, Lee SY. Regulation of Drosophila-virus interaction. Dev Comp Immunol. 2012;36: 262–266.

15. Öhlund P, Lundén H, Blomström A-L. Insect-specific virus evolution and potential effects on vector competence. Virus Genes. 2019;55: 127–137.

16. Bonning BC, Allen Miller W. Dicistroviruses. Annual Review of Entomology. 2010. pp. 129–150. doi: 10.1146/annurev-ento-112408-085457

17. Carreck NL, Ball BV, Martin SJ. Honey bee colony collapse and changes in viral prevalence associated with Varroa destructor. Journal of Apicultural Research. 2010. pp. 93–94. doi: 10.3896/ibra.1.49.1.13

18. Santillán-Galicia MT, Teresa Santillán-Galicia M, Ball BV, Clark SJ, Alderson PG. Transmission of deformed wing virus and slow paralysis virus to adult bees (Apis melliferaL.) byVarroa destructor. Journal of Apicultural Research. 2010. pp. 141–148. doi:10.3896/ibra.1.49.2.01

19. Stokstad E. Bee Virus Endemic. Science. 2007. p. 901b–901b. doi: 10.1126/science.318.5852.901b

20. Arnold PA, Johnson KN, White CR. Physiological and metabolic consequences of viral infection in Drosophila melanogaster. J Exp Biol. 2013;216: 3350–3357.

21. Cherry S, Perrimon N. Entry is a rate-limiting step for viral infection in a Drosophila melanogaster model of pathogenesis. Nature Immunology. 2004. pp. 81–87. doi: 10.1038/ni1019

22. Chtarbanova S, Lamiable O, Lee K-Z, Galiana D, Troxler L, Meignin C, et al. Drosophila C virus systemic infection leads to intestinal obstruction. J Virol. 2014;88: 14057–14069.

23. Blitvich BJ, Firth AE. Insect-Specific Flaviviruses: A Systematic Review of Their Discovery, Host Range, Mode of Transmission, Superinfection Exclusion Potential and Genomic Organization. Viruses. 2015;7: 1927–1959.

24. Li C-X, Shi M, Tian J-H, Lin X-D, Kang Y-J, Chen L-J, et al. Unprecedented genomic diversity of RNA viruses in arthropods reveals the ancestry of negative-sense RNA viruses. Elife. 2015;4. doi: 10.7554/eLife.05378

25. Shi M, Lin X-D, Tian J-H, Chen L-J, Chen X, Li C-X, et al. Redefining the invertebrate RNA virosphere. Nature. 2016. pp. 539–543. doi:10.1038/nature20167

26. Shi C, Zhao L, Atoni E, Zeng W, Hu X, Matthijnssens J, et al. The conservation of a core virome in Aedes mosquitoes across different developmental stages and continents. doi: 10.1101/2020.04.23.058701

27. Olmo RP, Martins NE, Aguiar ERGR, Marques JT, Imler J-L. The insect reservoir of biodiversity for viruses and for antiviral mechanisms. An Acad Bras Cienc. 2019;91 Suppl 3: e20190122.

28. Bronkhorst AW, van Rij RP. The long and short of antiviral defense: small RNA-based immunity in insects. Current Opinion in Virology. 2014. pp. 19–28. doi: 10.1016/j.coviro.2014.03.010

29. Gammon DB, Mello CC. RNA interference-mediated antiviral defense in insects. Curr Opin Insect Sci. 2015;8: 111–120.

30. van Rij RP, Saleh M-C, Berry B, Foo C, Houk A, Antoniewski C, et al. The RNA silencing endonuclease Argonaute 2 mediates specific antiviral immunity in Drosophila melanogaster. Genes Dev. 2006;20: 2985–2995.

31. Miesen P, Joosten J, van Rij RP. PIWIs Go Viral: Arbovirus-Derived piRNAs in Vector Mosquitoes. PLoS Pathog. 2016; 12: e1006017.

32. Whitfield ZJ, Dolan PT, Kunitomi M, Tassetto M, Seetin MG, Oh S, et al. The Diversity, Structure, and Function of Heritable Adaptive Immunity Sequences in the Aedes aegypti Genome. Current Biology. 2017. pp. 3511–3519.e7. doi: 10.1016/j.cub.2017.09.067

33. Tassetto M, Kunitomi M, Whitfield ZJ, Dolan PT, Sánchez-Vargas I, Garcia-Knight M, et al. Author response: Control of RNA viruses in mosquito cells through the acquisition of vDNA and endogenous viral elements. 2019. doi: 10.7554/elife.41244.026

34. Suzuki Y, Baidaliuk A, Miesen P, Frangeul L, Crist AB, Merkling SH, et al. Non-retroviral Endogenous Viral Element Limits Cognate Virus Replication in Aedes aegypti Ovaries. Current Biology. 2020. pp. 3495–3506.e6. doi: 10.1016/j.cub.2020.06.057

35. Vieira CB, Praça YR, da Silva Bentes KL, Santiago PB, Silva SMM, dos Santos Silva G, et al. Triatomines: Trypanosomatids, Bacteria, and Viruses Potential Vectors? Frontiers in Cellular and Infection Microbiology. 2018. doi: 10.3389/fcimb.2018.00405

36. Muscio OA, La Torre JL, Scodeller EA. Characterization of Triatoma virus, a picorna-like virus isolated from the triatomine bug Triatoma infestans. J Gen Virol. 1988;69 (Pt 11): 2929–2934.

37. Sánchez-Eugenia R, Méndez F, Querido JFB, Silva MS, Guérin DMA, Rodríguez JF. Triatoma virus structural polyprotein expression, processing and assembly into virus-like particles. J Gen Virol. 2015;96: 64–73.

38. Squires G, Pous J, Agirre J, Rozas-Dennis GS, Costabel MD, Marti GA, et al. Structure of the Triatoma viruscapsid. Acta Crystallographica Section D Biological Crystallography. 2013. pp. 1026–1037. doi:10.1107/s0907444913004617

39. Brito T, Julio A, Berni M, de Castro Poncio L, Bernardes ES, Araujo H, et al. Transcriptomic and functional analyses of the piRNA pathway in the Chagas disease vector Rhodnius prolixus. PLoS Negl Trop Dis. 2018; 12: e0006760.

40. Paim RMM, Araujo RN, Lehane MJ, Gontijo NF, Pereira MH. Long-term effects and parental RNAi in the blood feeder Rhodnius prolixus (Hemiptera; Reduviidae). Insect Biochemistry and Molecular Biology. 2013. pp. 1015–1020. doi:10.1016/j.ibmb.2013.08.008

41. Dong Y, Chao J, Liu J, Rice A, Holdbrook R, Liu Y, et al. Characterization of a novel RNA virus from Nesidiocoris tenuis related to members of the genus Iflavirus. Archives of Virology. 2018. pp. 571–574. doi:10.1007/s00705-017-3622-8

42. de Miranda JR, Dainat B, Locke B, Cordoni G, Berthoud H, Gauthier L, et al. Genetic characterization of slow bee paralysis virus of the honeybee (Apis mellifera L.). J Gen Virol. 2010;91: 2524–2530.

43. Pringle FM, Kalmakoff J, Ward VK. Analysis of the capsid processing strategy of Thosea asigna virus using baculovirus expression of virus-like particles. J Gen Virol. 2001;82: 259–266.

44. Sõmera M, Sarmiento C, Truve E. Overview on Sobemoviruses and a Proposal for the Creation of the Family Sobemoviridae. Viruses. 2015. pp. 3076–3115. doi:10.3390/v7062761

45. Büttner C, von Bargen S, Bandte M. Phytopathogenic Viruses. Principles of Plant-Microbe Interactions. 2015. pp. 115–122. doi: 10.1007/978-3-319-08575-3_13

46. Ribeiro JMC, Genta FA, Sorgine MHF, Logullo R, Mesquita RD, Paiva-Silva GO, et al. An insight into the transcriptome of the digestive tract of the bloodsucking bug, Rhodnius prolixus. PLoS Negl Trop Dis. 2014;8: e2594.

47. Brennecke J, Aravin AA, Stark A, Dus M, Kellis M, Sachidanandam R, et al. Discrete small RNA-generating loci as master regulators of transposon activity in Drosophila. Cell. 2007; 128: 1089–1103.

48. Gunawardane LS, Saito K, Nishida KM, Miyoshi K, Kawamura Y, Nagami T, et al. A slicer-mediated mechanism for repeat-associated siRNA 5’ end formation in Drosophila. Science. 2007;315: 1587–1590.

49. Ciota AT, Bialosuknia SM, Ehrbar DJ, Kramer LD. Vertical Transmission of Zika Virus by Aedes aegypti and Ae. albopictus Mosquitoes. Emerg Infect Dis. 2017;23: 880–882.

50. Ferreira-de-Lima VH, Lima-Camara TN. Natural vertical transmission of dengue virus in Aedes aegypti and Aedes albopictus: a systematic review. Parasit Vectors. 2018; 11: 77.

51. Thangamani S, Huang J, Hart CE, Guzman H, Tesh RB. Vertical Transmission of Zika Virus in Aedes aegypti Mosquitoes. The American Journal of Tropical Medicine and Hygiene. 2016. pp. 1169–1173. doi: 10.4269/ajtmh.16-0448

52. Santos D, Mingels L, Vogel E, Wang L, Christiaens O, Cappelle K, et al. Generation of Virus- and dsRNA-Derived siRNAs with Species-Dependent Length in Insects. Viruses. 2019; 11. doi: 10.3390/v11080738

53. Goic B, Vodovar N, Mondotte JA, Monot C, Frangeul L, Blanc H, et al. RNA-mediated interference and reverse transcription control the persistence of RNA viruses in the insect model Drosophila. Nat Immunol. 2013; 14: 396–403.

54. Lan H, Wang H, Chen Q, Chen H, Jia D, Mao Q, et al. Small interfering RNA pathway modulates persistent infection of a plant virus in its insect vector. Sci Rep. 2016;6: 20699.

55. Fernández-Presas AM, Padilla-Noriega L, Becker I, Robert L, Jiménez JA, Solano S, et al. Enveloped and non-enveloped viral-like particles in Trypanosoma cruzi epimastigotes. Rev Inst Med Trop Sao Paulo. 2017;59: e46.

56. Pane A, Jiang P, Zhao DY, Singh M, Schüpbach T. The Cutoff protein regulates piRNA cluster expression and piRNA production in the Drosophila germline. EMBO J. 2011;30: 4601–4615.

57. Pritykin Y, Brito T, Schupbach T, Singh M, Pane A. Integrative analysis unveils new functions for the Cutoff protein in noncoding RNA biogenesis and gene regulation. RNA. 2017;23: 1097–1109.

58. Mesquita RD, Vionette-Amaral RJ, Lowenberger C, Rivera-Pomar R, Monteiro FA, Minx P, et al. Genome of Rhodnius prolixus, an insect vector of Chagas disease, reveals unique adaptations to hematophagy and parasite infection. Proc Natl Acad Sci U S A. 2015; 112: 14936–14941.

59. Dobin A, Davis CA, Schlesinger F, Drenkow J, Zaleski C, Jha S, et al. STAR: ultrafast universal RNA-seq aligner. Bioinformatics. 2013;29: 15–21.

60. Haas BJ, Papanicolaou A, Yassour M, Grabherr M, Blood PD, Bowden J, et al. De novo transcript sequence reconstruction from RNA-seq using the Trinity platform for reference generation and analysis. Nat Protoc. 2013;8: 1494–1512.

61. Patro R, Duggal G, Love MI, Irizarry RA, Kingsford C. Salmon provides fast and bias-aware quantification of transcript expression. Nat Methods. 2017; 14: 417–419.

62. Li W, Jaroszewski L, Godzik A. Clustering of highly homologous sequences to reduce the size of large protein databases. Bioinformatics. 2001. pp. 282–283. doi: 10.1093/bioinformatics/17.3.282

63. Li W, Jaroszewski L, Godzik A. Tolerating some redundancy significantly speeds up clustering of large protein databases. Bioinformatics. 2002; 18: 77–82.

64. Smith-Unna R, Boursnell C, Patro R, Hibberd JM, Kelly S. TransRate: reference-free quality assessment of de novo transcriptome assemblies. Genome Res. 2016;26: 1134–1144.

65. UniProt Consortium. UniProt: a worldwide hub of protein knowledge. Nucleic Acids Res. 2019;47: D506–D515.

66. Song L, Shankar DS, Florea L. Rascaf: Improving Genome Assembly with RNA Sequencing Data. Plant Genome. 2016;9. doi: 10.3835/plantgenome2016.03.0027

67. Altschul SF, Gish W, Miller W, Myers EW, Lipman DJ. Basic local alignment search tool. J Mol Biol. 1990;215: 403–410.

68. Bushmanova E, Antipov D, Lapidus A, Prjibelski AD. rnaSPAdes: a de novo transcriptome assembler and its application to RNA-Seq data. Gigascience. 2019;8. doi: 10.1093/gigascience/giz100

69. Marchler-Bauer A, Anderson JB, Cherukuri PF, DeWeese-Scott C, Geer LY, Gwadz M, et al. CDD: a Conserved Domain Database for protein classification. Nucleic Acids Res. 2005;33: D192–6.

70. Thompson JD, Higgins DG, Gibson TJ. CLUSTAL W: improving the sensitivity of progressive multiple sequence alignment through sequence weighting, position-specific gap penalties and weight matrix choice. Nucleic Acids Res. 1994;22: 4673–4680.

71. Kumar S, Stecher G, Li M, Knyaz C, Tamura K. MEGA X: Molecular Evolutionary Genetics Analysis across Computing Platforms. Mol Biol Evol. 2018;35: 1547–1549.

72. The neighbor-joining method: a new method for reconstructing phylogenetic trees. Molecular Biology and Evolution. 1987. doi: 10.1093/oxfordjournals.molbev.a040454

73. Felsenstein J. CONFIDENCE LIMITS ON PHYLOGENIES: AN APPROACH USING THE BOOTSTRAP. Evolution. 1985;39: 783–791.

74. Jones DT, Taylor WR, Thornton JM. The rapid generation of mutation data matrices from protein sequences. Bioinformatics. 1992. pp. 275–282. doi: 10.1093/bioinformatics/8.3.275

75. Zhang K-Y, Gao Y-Z, Du M-Z, Liu S, Dong C, Guo F-B. Vgas: A Viral Genome Annotation System. Front Microbiol. 2019; 10: 184.

76. Ramírez F, Ryan DP, Grüning B, Bhardwaj V, Kilpert F, Richter AS, et al. deepTools2: a next generation web server for deep-sequencing data analysis. Nucleic Acids Res. 2016;44: W160–5.

77. Kent WJ. The Human Genome Browser at UCSC. Genome Research. 2002. pp. 996–1006. doi: 10.1101/gr.229102.

78. Langmead B, Trapnell C, Pop M, Salzberg SL. Ultrafast and memory-efficient alignment of short DNA sequences to the human genome. Genome Biol. 2009; 10: R25.

79. Quinlan AR. BEDTools: The Swiss-Army Tool for Genome Feature Analysis. Current Protocols in Bioinformatics. 2014. pp. 11.12.1–11.12.34. doi: 10.1002/0471250953.bi1112s47

80. Rice P, Longden I, Bleasby A. EMBOSS: The European Molecular Biology Open Software Suite. Trends in Genetics. 2000. pp. 276–277. doi: 10.1016/s0168-9525(00)02024-2

81. Giraldo-Calderón GI, Emrich SJ, MacCallum RM, Maslen G, Dialynas E, Topalis P, et al. VectorBase: an updated bioinformatics resource for invertebrate vectors and other organisms related with human diseases. Nucleic Acids Res. 2015;43: D707–13.

